# Upper body and ankle strategies compensate for reduced lateral stability at very slow walking speeds

**DOI:** 10.1101/2020.07.16.207092

**Authors:** Aaron N. Best, Amy R. Wu

## Abstract

At the typical walking speeds of healthy humans, step placement seems to be the primary strategy to maintain gait stability, with ankle torques and upper body momentum providing additional compensation. The average walking speeds of populations with an increased risk of falling, however, are much slower and may require differing control strategies. The purpose of this study was to analyze mediolateral gait stability and the contributions of the different control strategies at very slow walking speeds. We analyzed an open dataset including kinematics and kinetics from eight healthy subjects walking at speeds from 0.1 to 0.6 m/s as well as a self-selected speed. As gait speed slowed, we found that the margin of stability decreased linearly. Increased lateral excursions of the extrapolated centre of mass, caused by increased lateral excursions of the trunk, were not compensated for by an equivalent increase in the lateral centre of pressure, leading to decreased margin of stability. Additionally, both the ankle eversion torque and hip abduction torque at the minimum margin of stability event increased at the same rate as gait speed slowed. These results suggest that the contributions of both the ankle and the upper body to stability are more crucial than stepping at slow speeds, which have important implications for populations with slow gait and limited motor function.

## Introduction

In populations with increased fall risk, the typical walking speeds may be less than half of the walking speeds of healthy humans [1, 2, 3]. Slower walking speeds are associated with decreased quality of life, increased hospitalization and risk of death [4, 5]. One common cause of falls in elderly populations and other clinical groups is the improper shifting of weight during forward walking [6, 7, 8]. Elucidation of the factors that lead to instability for these populations necessitates investigations into both the active control and passive mechanisms that enable healthy humans to maintain gait stability across a range of walking speeds. Much research in gait stability mechanisms, however, is conducted at the typical walking speeds of healthy humans, from 0.6 m/s to 2.0 m/s [9, 10, 11].

Stable gait can be achieved through control of the body centre of mass (CoM) relative to the base of support (BoS) created by contact between the feet and the ground [12]. This control entails passive and active mechanisms that could involve both the upper and lower body. While the sagittal plane can be passively stabilized, lateral balance requires active stabilization [13]. Lower limb control consists of the stepping strategy and the ankle strategy [14]. Stepping is thought to be the most important strategy [15], as it can compensate for larger perturbations than the ankle strategy through corrective foot placement [14]. The ankle strategy is used to modulate the centre of pressure (CoP) under the stance foot. Although the size of the foot limits the maximum distance the CoP can travel, the ankle strategy is faster than the stepping strategy, as it can be applied before the next foot placement [14, 16]. The upper body, or trunk, can also be used to maintain stability [17]. Rotation of the trunk about the body CoM impacts the acceleration of the CoM [18], allowing for the trunk to control the position of the CoM and redirect it towards the interior of the BoS [19]. The trunk strategy has been observed in scenarios where stepping strategy is confined [19] or an unfavourable strategy [20].

Previous investigations on the impact of gait speed on CoM behaviour suggest active control could be required. Using Lyapunov exponents to evaluate stability between 0.62 m/s and 1.72 m/s, Bruijn et al. [10] found that the long-term local divergence exponent suggested a decrease in stability. However, their overall results were inconclusive as they also found that the short-term exponent, which is perhaps more closely related to stability [21], suggested an increase in stability. Orendurff et al. [22] investigated the frontal plane motion of the body CoM from subjects walking between 0.7 m/s and 1.6 m/s.They found that, as gait speed decreased, the lateral displacement of the CoM increased. The greater displacement reduces the lateral distance between the CoM and the BoS and thus may reduce stability without some compensation from the body [12]. A possible continued increase in CoM displacement at speeds slower than 0.7 m/s could further impact gait stability.

One potential strategy to compensate for the increase in lateral CoM displacement would be to take wider steps and maintain a constant or greater distance between the CoM and the BoS, yet step width modulation does not seem well supported at slow walking speeds. A study by Stimpson et al. [23] suggests that stepping might contribute less to gait stability at slower speeds. They tested walking speeds from 0.2 to 1.2 m/s and found that the relationship between pelvis state and step-width weakened as speed slowed. Wu et al. [24] investigated speeds as low as 0.1 m/s and found that as gait speed slowed, the percentage of gait spent in stance increased. This increase may lead to a reduction in the ability to use the stepping strategy as the foot spends more time in contact with the ground. The same study also found no significant changes in step width with walking speed. If the step-width remains constant, then the distance between the CoM and the BoS will decrease as lateral CoM position increases, necessitating additional stability mechanisms.

Perhaps, then an ankle or trunk strategy or both could provide the necessary compensation to maintain gait stability at slow speeds. den Otter et al. [25] investigated muscle activation of the lower limb muscles with surface electromyography at gait speeds from 0.06 to 1.39 m/s. As speed slowed, the activation of the peroneus longus, which applies an eversion moment about the ankle joint, increased. The greater activation suggests that the ankle strategy may provide additional compensation at slower speeds. However, this study did not measure the activity of upper body muscles, and the possibility of increased contribution from the trunk is unknown.

Here, we investigated mediolateral gait stability at very slow walking speeds, including the contributions of the upper and lower body stabilization strategies. We used an open dataset [24] which had previously concluded that step-width remained constant and the stance time increased as gait speed slowed. We hypothesized that as gait speed slows, the body CoM excursions would increase, causing a lower margin of stability (MoS) as the step-width remained constant. Additionally, with the longer stance times, we expected that the stepping strategy would become less dominant, and the lateral ankle strategy would provide more compensation for stability to be maintained.

## Methods

Kinematic and kinetic data from an existing slow walking dataset was used to test our hypotheses [24]. The dataset contained normative data for eight healthy subjects (six female, two male, weight 65.6 *±* 9.62 kg (mean *±* s.d.), age 23 to 31 years) walking at 0.1, 0.3, 0.5, and 0.6 m/s as well as a self-selected speed (ranging from 0.92 to 1.14 m/s). Randomized walking conditions were performed for two minutes while walking on an instrumented treadmill with optical motion capture. The data used was stride-by-stride normalized kinematics of the body CoM, trunk CoM, left and right CoP as well as the normalized hip and ankle torques in the frontal plane. The positions and torques were calculated using standard kinematic and inverse dynamics in OpenSim [26] with procedures outlined in the previous study [24].

To quantify gait stability, we calculated the average minimum MoS during single leg support for each speed. The MoS was computed as the mediolateral distance from the CoP of the stance leg to the body extrapolated centre of mass (XCoM) [12]. The XCoM is defined as:

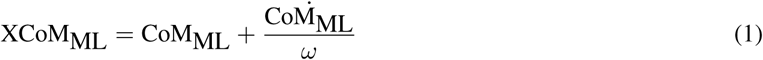

where 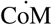 is the CoM velocity and *ω* is the first eigenfrequency of the inverted pendulum model of walking. The pendulum length used in the eigenfrequency estimation was the magnitude of the vector from the CoP of the stance leg to body CoM.

To further investigate the relationship between body kinematics and stability with gait speed, the state of the upper and lower body at the minimum MoS was analyzed. Specifically, the trunk and CoP positions were used to evaluate the contribution of the upper and lower body to the change in the MoS. This behavior was averaged between the left and right minimum MoS events. Mediolateral hip and ankle torques were also analyzed at the minimum MoS and throughout the gait cycle. In the analysis of the hip torques, only six subjects were used as the hip torque curves for two of the subjects were found to be irregular.

Speed-related changes were quantified using linear fits between the measures and gait speed. For each linear fit, there was one common slope with individual offsets for each subject. Statistics were performed on the regression trend coefficients using t-statistics with a significance level of *α* = 0.05. All reported values have been normalized with a combination of leg length *L*, mass *M* and gravity *g*. Length was divided by *L* (0.91 *±* 0.04 m), velocity by 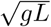, and acceleration by *g*. Torque was divided by *MgL* (587 *±* 108 N-m), and time was divided by 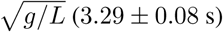

## Results

We found that the MoS decreased as gait speed slowed. Along with XCoM, both the lateral excursion of the trunk CoM and body CoM increased with slower speeds. The CoP excursion at the minimum MoS event also increased but could not match the XCoM’s rate of increase, leading to lower margins. At the same minimum MoS event, the ankle eversion torque and hip abduction torque also grew in magnitude as gait speed slowed.

As gait speed varied, the kinematics of the body and trunk CoM varied while the BoS created by the CoP remained relatively constant (Figure 1A). The trunk CoM displayed larger lateral excursions as gait speed slowed. Similarly, the range of the body CoM excursions increased at 0.07 m/ms^−1^ (Figure 1B and Table 1). Despite changes in gait speed, the mediolateral CoM velocity remained similar across speed conditions (Figure 1B and Table 1). As the amplitude of the CoM motion increased and the velocity remained constant, the mediolateral acceleration of the CoM decreased as speed slowed (Figure 1B and Table 1).

**Table 1:** Summary of the regression results and statistics. All slopes (mean *±* 95% confidence interval, CI) and offsets (mean *±* s.d.) are in reported in dimensionless units. *R*^2^ values indicate the goodness of fit for the linear regressions. The asterisk indicates a statistically significant trend (^*\**^*p <* 0.05).

| Lateral Measure | Slope (Mean ± CI) | $p$-value | Offset (Mean ± s.d.) | $R^2$ |
| --- | --- | --- | --- | --- |
| CoM Position Range | $-0.24 \pm 0.03$ | $1.0e-15^*$ | $0.13 \pm 0.01$ | 0.91 |
| CoM Velocity Range | $0.05 \pm 0.05$ | 0.052 | $0.10 \pm 0.02$ | 0.63 |
| CoM Acceleration Range | $0.50 \pm 0.07$ | $3.3e-16^*$ | $0.06 \pm 0.02$ | 0.91 |
| MoS | $0.07 \pm 0.01$ | $5.1e-12^*$ | $0.005 \pm 0.004$ | 0.84 |
| CoP | $-0.05 \pm 0.03$ | $3.6e-3^*$ | $0.10 \pm 0.01$ | 0.81 |
| XCoM | $-0.07 \pm 0.02$ | $2.2e-10^*$ | $0.079 \pm 0.005$ | 0.91 |
| Trunk CoM | $-0.16 \pm 0.03$ | $9.5e-12^*$ | $0.07 \pm 0.01$ | 0.81 |
| Ankle Torque | $0.036 \pm 0.008$ | $3.7e-11^*$ | $-0.019 \pm 0.002$ | 0.91 |
| Hip Torque | $0.036 \pm 0.02$ | $1.1e-3^*$ | $-0.080 \pm 0.006$ | 0.72 |

**Figure 1:**
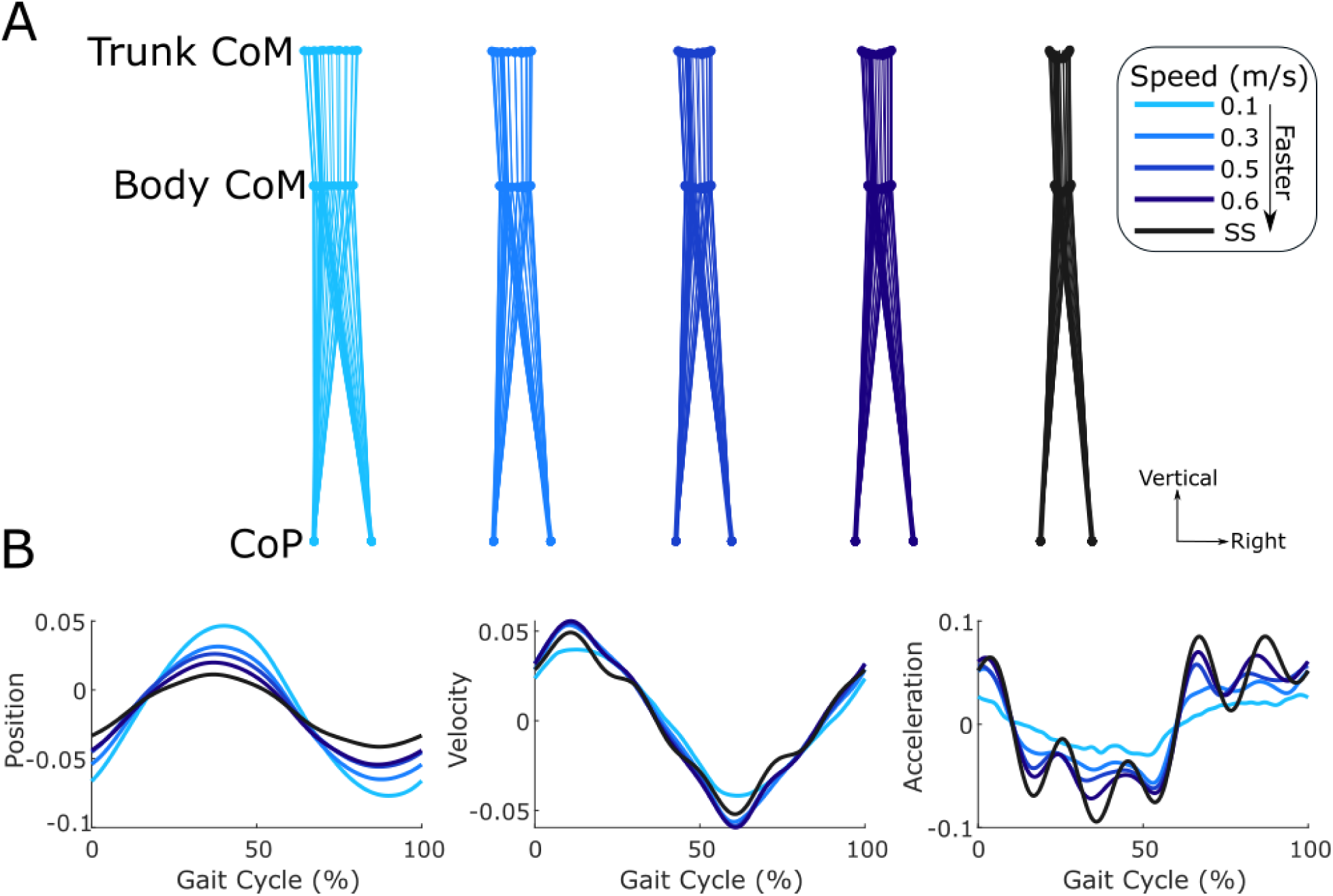
Average mediolateral gait kinematics for all subjects (*N* = 8) across the different gait speed conditions. (A) Average trajectories of the body CoM and trunk CoM throughout the gait cycle and the left and right CoP at right heel strike. The range of motion of the trunk and body CoM decreased with gait speed, but the CoP remained relatively constant. (B) Average mediolateral body CoM position, velocity and acceleration. CoM position moved more laterally as gait speed slowed, accompanied by relatively constant CoM velocity magnitudes and decreased CoM accelerations.

Slower gait speeds induced changes in XCoM and CoP behavior, leading to a decrease in minimum MoS (Figure 2, Table 1). Qualitatively, the trajectories of the XCoM, CoM, and CoP appeared similar across the different gait speeds with a smaller percentage of the gait cycle spent in single support at slower speeds (Figure 2A). Despite the changes in timing, the minimum MoS consistently occurred after the transition from double to single support. As gait speed decreased, the average minimum MoS decreased linearly (Figure 2B and Table 1) at a rate of 0.02 m/ms^−1^.

**Figure 2:**
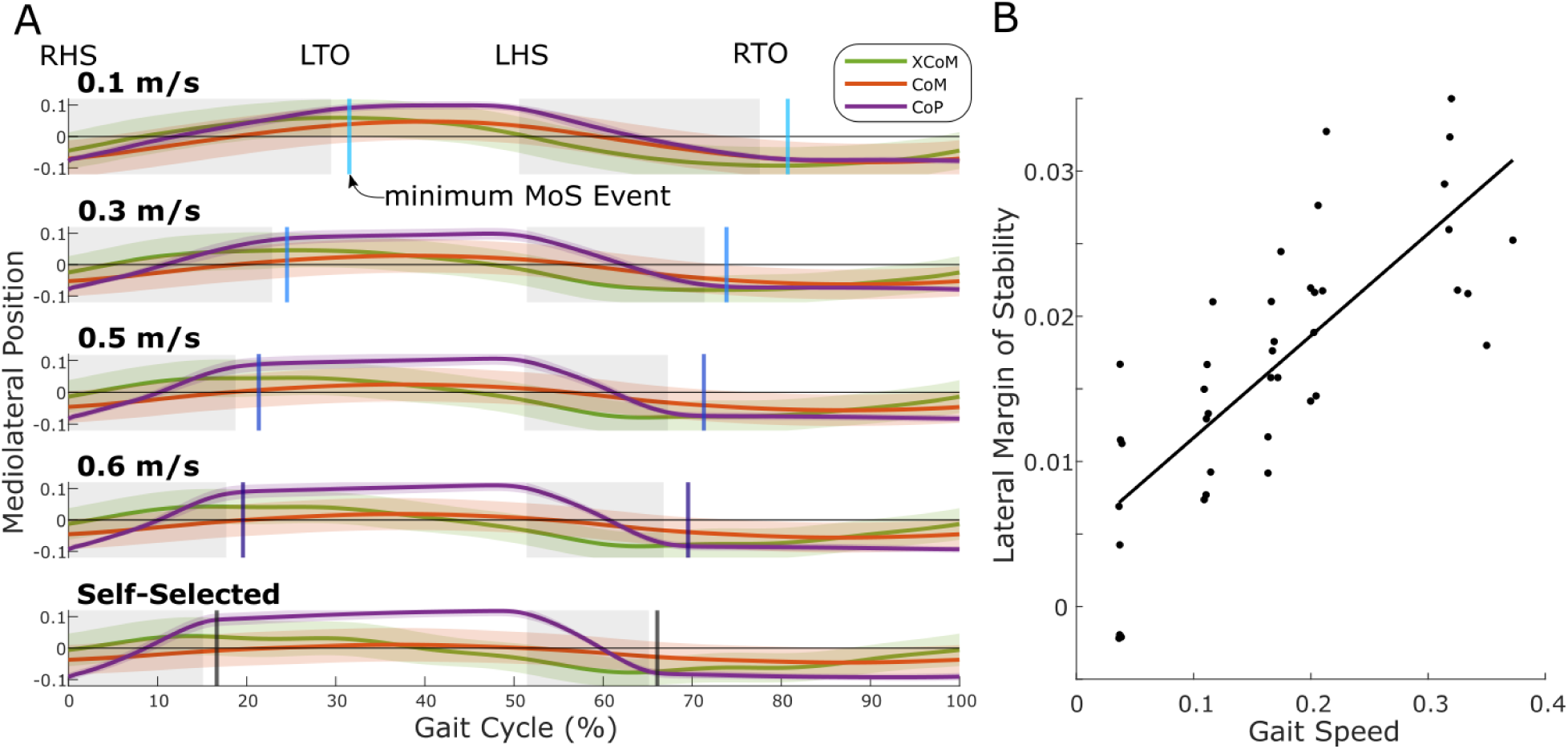
Mediolateral MoS and its contributors across the different gait speed conditions (*N* = 8). (A) Average trajectories of the total CoP, body CoM and XCoM. The average minimum MoS (vertical lines) consistently occurs shortly after the transition from double support (shaded region) to single support (unshaded region). The average location of gait events are indicated (RHS: right heelstrike, LTO: left toe off, LHS: left heelstrike, and RTO: right toe off). The percentage of the gait cycle spent in single support is shorter at slower speeds. (B) Linear regression of the minimum MoS. The average minimum MoS of individual subjects (dots) and its corresponding linear fit (solid line) demonstrate a significant decrease with slower gait speeds (*p <* 0.05).

At the minimum MoS event, the CoP, XCoM and trunk CoM were all more lateral at slower gait speeds (Figure 3 and Table 1). However, the rate at which the XCoM moved laterally was 1.4 times greater than the rate at which the CoP moved laterally. The lateral position of the trunk decreased the most drastically, decreasing at a rate of 0.049 m/ms^−1^.

**Figure 3:**
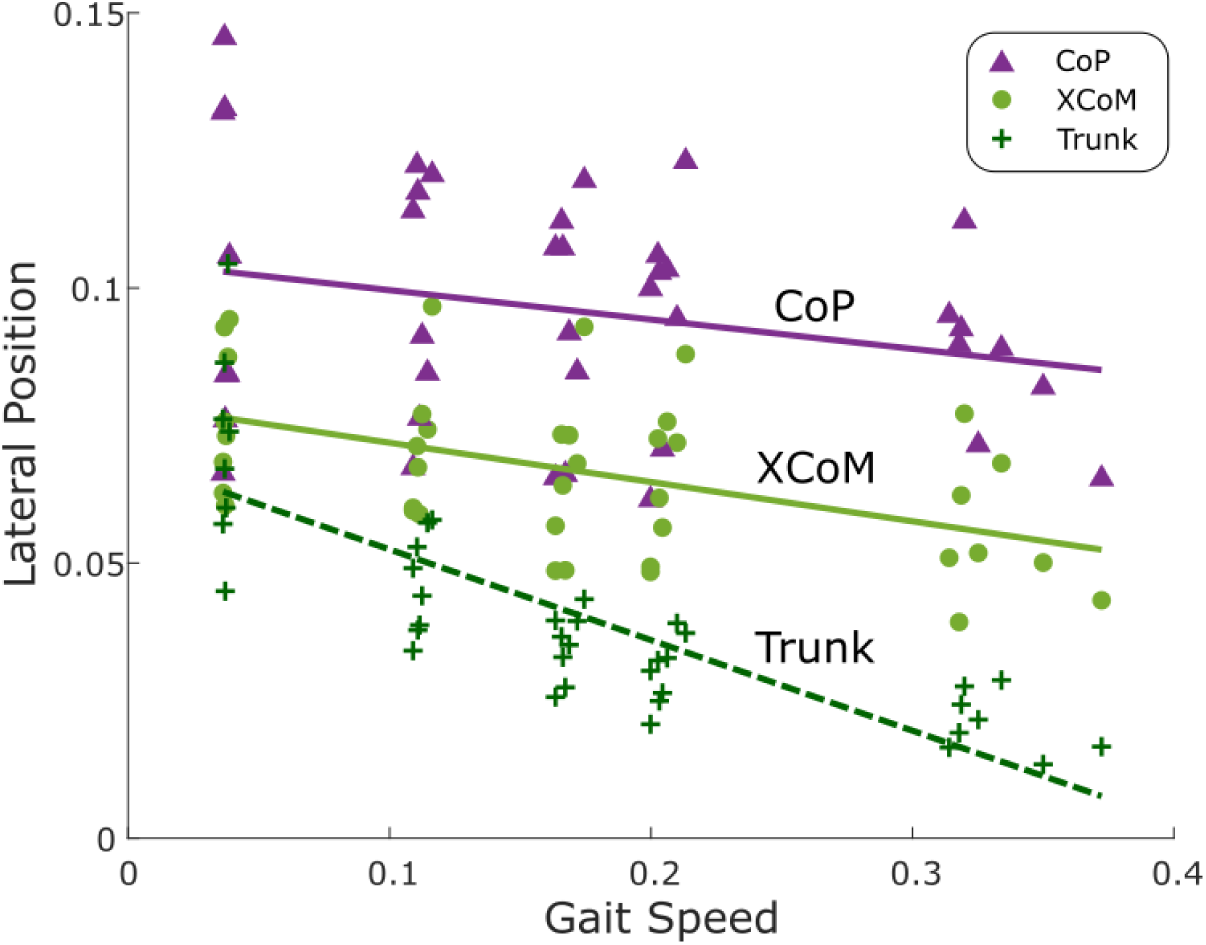
Linear regression of the lateral position of the CoP, XCoM and trunk CoM at the minimum MoS event. The average lateral position of the CoP, XCoM or trunk CoM at the minimum MoS event for each individual subject (dots) and their fits (solid and dashed lines) are shown (*N* = 8). Both the lateral CoP and XCoM position increased with slower gait speed at the minimum MoS (*p <* 0.05). The rate at which the XCoM lateral position increased was faster than that of the CoP. The lateral trunk CoM position, which affects the body CoM and XCoM, also increased as gait speed decreased (*p <* 0.05).

Mediolateral ankle and hip torque were both affected by walking speed, with increased contributions at the minimum MoS event. Qualitatively, the ankle torque profiles were fairly consistent across gait speeds with lower peak eversion torques at slower speeds (Figure 4A top). Similarly, the hip torques were similar among the different gait speed conditions but with reduced peak magnitudes as well (Figure 4A bottom). At the minimum MoS event, both the ankle eversion torque and the hip abduction torque increased in magnitude as gait speed decreased (Figure 4B and Table 1). The rate at which the torques increased was similar for both at 7.09 N-m/ms^−1^ (Table 1), representing average torque increase of 2.5 times and 1.2 times from the slowest speed and the self-selected (fastest) speed for the ankle and hip respectively.

**Figure 4:**
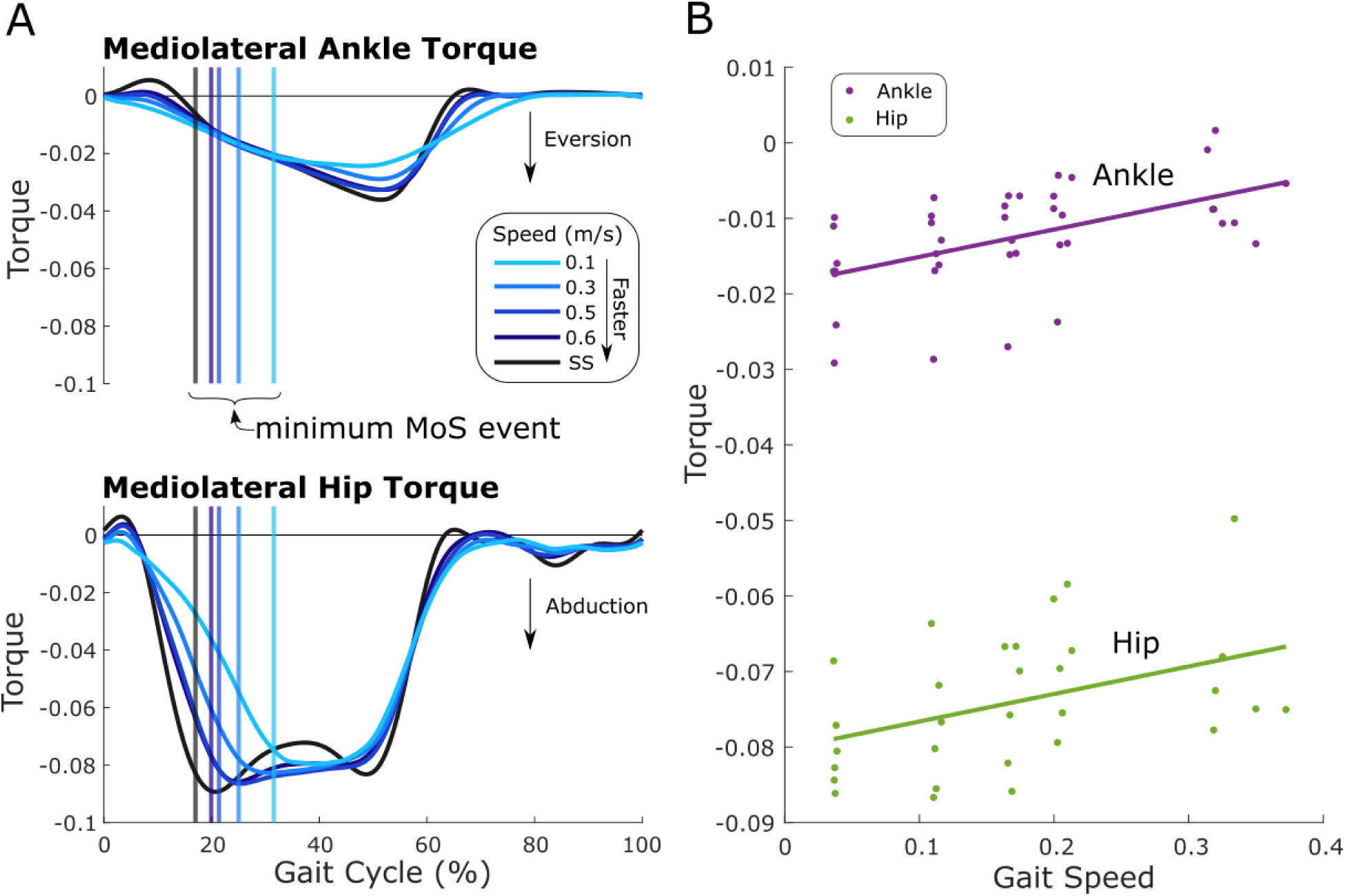
Mediolateral ankle and hip torques across different gait speeds. (A) The average right hip (*N* = 6) and ankle (*N* = 8) torques as a function of the gait cycle. The average location of the minimum MoS of the right leg for the respective gait speed is indicated by vertical lines. Qualitatively, the ankle and hip trajectories are similar across the speeds with lowered peak eversion and abduction torques, respectively, for slower speeds.(B) Linear regression of the mediolateral ankle and hip torques at the minimum MoS. The average torque of the right and left ankle (*N* = 8) and hip (*N* = 6) at the minimum MoS event for each individual subject (dots) are shown along with their fits (solid lines). As gait speed slowed, both the hip abduction torque and the ankle eversion torque increased in magnitude (*p <* 0.05).

## Discussion

In the current study, we investigated the effects of gait speed on mediolateral gait stability at very slow speeds. We hypothesized that as gait speed decreased, the MoS would decrease. Our results agreed with this hypothesis, finding that the MoS decreased linearly as gait speed slowed (Figure 2B). As speed decreased, the amplitude of the trunk CoM increased, causing greater excursions of the XCoM that were not compensated by an equivalent increase in the lateral CoP. We also expected that as gait speed slowed, stance time increased, and step width remained unchanged, the ankle strategy would become more dominant. We found that the ankle eversion torque increased at the minimum MoS event, along with the hip abduction torque (Figure 4B). These results suggest that the ankle and hip strategy are needed to maintain stability at slower speeds when stepping is no longer used or effective.

Maintaining gait stability is crucial, but energetic efficiency also governs walking behavior [27, 28, 29]. These two determinants of gait may conflict, possibly leading to trade-offs where gait stability is reduced in order to achieve a more efficient gait. As gait speed decreased, the mediolateral CoM velocity remained relatively constant while CoM excursion increased, allowing CoM acceleration to decrease (Figure 1B). This behavior seems disadvantageous for gait stability as the increased excursions of the CoM led to a lower MoS at slower speeds (Figure 2B). If instead the subjects had elected to decrease their mediolateral CoM velocity and maintain a constant range of CoM excursion, yielding the same decrease in the CoM acceleration, then the MoS could increase. The position term in the XCoM (Equation 1) would remain constant, but the velocity term would decrease, reducing the range of the XCoM. Despite the possible benefits of the latter option, subjects elected to maintain a relatively constant mediolateral CoM velocity at the cost of reduced MoS due to greater CoM amplitudes. This preferred but seemingly unfavourable behaviour of increasing CoM amplitude may be the result of a trade-off between gait stability and energetic costs. Previous studies have suggested such trade-offs, where the elected step-width avoids loss of balance but is not overly energetically costly [23, 30]. It is also possible that there may be an associated energetic cost with decreases in the CoM motion as it would require the motion of the trunk to be restricted. Further exploration of the potential energetic costs of reducing CoM velocity requires additional experimentation.

Our finding of decreasing MoS with reducing gait speed (Figure 2) is in agreement with previous studies that investigated gait in a similar range of speeds. Stimpson et al. [23] also found that the MoS generally decreased as gait speed decreased. However, they observed a significant increase in the MoS from 0.4 m/s to 0.2 m/s that was not observed in the current study. The difference may arise from the time in the gait cycle where the MoS was calculated. We calculated the minimum MoS during single support of each leg whereas the previous study calculated the MoS at the end of each step. Additionally, Stimpson et al. found that step-width increased at slower speeds, which is contrary to the previously published results of the dataset [24] we used here. The relationship between step-width and gait speed remains inconclusive as some studies, including the current study, found no significant difference [31, 24] and others have found a significant difference [22, 23].

We had hypothesized that the lateral ankle strategy would provide compensation for the reduced contribution of the stepping strategy based on the previously observed increase in muscle activation of the peroneus longus [25]. In agreement with our hypothesis, we found that the ankle eversion torque increased as gait speed slowed (Figure 4B). The increased activity of the peroneus longus muscle, which acts to evert the ankle, was thought to prevent excessive ankle inversion during single support [25], caused by the increased medial ground reaction force at slower speeds [32]. Yet, the medial ground reaction forces decreased at slower speeds for this dataset [24]. These inconsistent findings suggest that the increased ankle eversion torque does not necessarily serve to counter an increased ground reaction force but instead could possibly control the location of the CoP within the BoS or redirect the ground reaction force. As gait speed slowed, the abduction hip torque also increased at the same rate as the eversion ankle torque (Figure 4B and Table 1). The hip torque could be responsible for the increasing lateral position of the CoP at the minimum MoS (Figure 3) or possibly stabilizing the trunk as it moved more laterally at slower speeds. The upper body behavior at slow speeds could be similar to that of balancing on one leg [19], where the hip moment varies the mediolateral component of the ground reaction force to bring a displaced CoM back over the BoS. A previous investigation of mediolateral balance in elderly adults has found that subjects with weaker hip abductor muscles displayed poorer performance in balance related tasks [33].

The current study has some limitations to be considered. The MoS was used as the primary measure of gait stability as its calculation accounts for both position and velocity of the body CoM. However, this simplistic measure is based on the inverted pendulum model of walking and does not consider balance strategies that utilize trunk momentum or hip torques. Other stability measures that may lead to further insights, such as local divergence exponents [34] or foot placement regressions [35], requires at least 150 strides [36] and could not be used due to the limited number of strides available in the existing dataset. These additional analyses would have provided further insights on the nature of the motion of the CoM as well as possible alterations to stepping control strategies at very slow speeds. Our analysis primarily focused on the minimum MoS event as the instance in which gait stability was the weakest and requiring more control. However, gait behaviour during other parts of stride is likely to also contribute towards stability. We also evaluated stability without external perturbations, which would have allowed for more thorough identification of compensatory reactions to destabilization events. Additionally, we reported the average behavior between the left and right legs but found some asymmetric behavior between the two legs that was not further analyzed.

Our results provide some interesting implications and directions for future work. Gait stability seems to decline as speed decreases, which could lead to an elevated risk of falls for populations with slow gait. The increase in trunk motion across gait speeds, accompanied by greater hip torques, signify that the upper body influences gait stability. The increase in ankle torque at the same rate as the hip suggests that the ankle strategy contributes as equally to lateral stabilization. It remains unclear whether the trunk motion arises from passive pendular mechanics or active control to stabilize gait. The CoM behaviour adopted at slow speeds also suggests that reduced margins in stability were preferred over, perhaps, the energetic cost of restricting CoM motion. Future work should be directed towards understanding the role of upper body control in the energetics and stability of gait.

## Ethics

The data used in this study was acquired from a publicly available dataset.

## Data Accessibility

The data used in the current study is available in Queen’s University Dataverse repository at https://doi.org/10.5683/SP2/EMQLLE.

## Authors’ Contribution

A.N.B and A.R.W conceived the study and drafted the manuscript. A.N.B conducted the data analysis. A.R.W acquired funding and supervised the study. A.N.B and A.R.W read and approved of the final manuscript.

## Acknowledgement

The authors would like to thank Edwin van Asseldonk and Mark Vlutters for their comments on the early results of the study.

